# Comparing DNA, RNA and protein levels for measuring microbial activity in nitrogen-amended soils

**DOI:** 10.1101/466680

**Authors:** Luis H. Orellana, Janet K. Hatt, Ramsunder Iyer, Karuna Chourey, Robert L. Hettich, Jim C. Spain, Wendy H. Yang, Joanne C. Chee-Sanford, Robert A. Sanford, Frank E. Löffler, Konstantinos T. Konstantinidis

**Author notes:** Address correspondence to Konstantinos T. Konstantinidis.

## Abstract

Multi-omic techniques can offer a comprehensive overview of microbial communities at the gene, transcript and protein levels. However, to what extent these levels reflect *in situ* process rates is less clear, especially in highly complex habitats such as soils. Here we performed microcosm incubations using soil from a site with a history of agricultural management. Microcosms, amended with isotopically labelled ammonium and urea to simulate a fertilization event, showed nitrification (up to 4.1 ± 0.87 µg N-NO_3_^-^ g^-1^ dry soil d^-1^) and accumulation of N_2_O after 192 hours of incubation. Nitrification activity (NH_4_^+^→NH_2_OH→NO_2_^-^→NO_3_^-^) was accompanied by a 6-fold increase in relative expression of the 16S rRNA gene (RNA/DNA) between 10 and 192 hours of incubation for ammonia-oxidizing bacteria (AOB) *Nitrosomonas* and *Nitrosospira*. In contrast, ammonia-oxidizing archaea (AOA) and complete ammonia oxidizer (comammox) nitrifiers showed stable gene expression during incubations but were generally more abundant (DNA level) than their *Betaproteobacteria* AOB counterparts. A strong relationship between nitrification activity and (mostly) betaproteobacterial ammonia monooxygenase (*amoA*; NH_4_^+^→NH_2_OH) and nitrite oxidoreductase (*nxrA*; NO_2_^-^→NO_3_^-^) transcript abundances revealed that mRNA levels quantitatively reflected measured activity and were generally more sensitive than the DNA level in the microcosm incubations. Although peptides related to housekeeping proteins from nitrite-oxidizing microorganisms were detected, their abundance was not significantly correlated with activity, revealing that meta-proteomics provided only a qualitative assessment of activity. Altogether, these findings underscore the strengths and limitations of multi-omic approaches for assessing complex microbial communities and provide the molecular means to assess nitrification processes in soils.

**IMPORTANCE:** Even though the use of omic approaches has expanded our knowledge of the diversity of microbial communities in natural and engineered systems, it is less clear how well the use of whole community DNA-, RNA- or protein-based approaches reflect microbial activities. To this end, we directly compared the different levels of molecular information (i.e., DNA, RNA or proteins) in order to assess which level best correlated with isotope-based measurements of nitrification activity in agricultural soils after fertilization. This work reveals the strengths as well as the associated limitations of metagenomic, metatranscriptomic, and metaproteomic approaches in serving as reliable proxies for examining microbial activities in highly diverse environments like soils.

## INTRODUCTION

Even though the central role of microbes in the cycling of nitrogen is recognized, the dynamics and controls of the interrelated microbial nitrogen pathways in agricultural soils are still poorly understood. This scarcity of information limits the development of more accurate, predictive models of nitrogen flux that encompasses the role of microbes in the generation and consumption of nitrogen substrates, as well as the emission of greenhouse gases, including nitrous oxide (N_2_O) (1). In agricultural soils receiving large inputs of nitrogen fertilizer, ammonia-oxidizing bacteria (AOB), ammonia-oxidizing archaea (AOA) and nitrite-oxidizing bacteria (NOB) collectively are responsible for the conversion of ammonium to nitrate. In addition, the recent discovery of *Nitrospira* bacteria capable of complete oxidation of ammonia to nitrate (comammox) has revealed that the process of nitrification in natural environments might be carried out by a single taxon (2, 3). It has also been reported that nitrification is a major N_2_O source under low oxygen concentrations (4), although detailed mechanistic understanding is lacking (5). Alternatively, under anoxic conditions, nitrate (NO_3_^-^) can be reduced to gaseous forms such as dinitrogen (N_2_), nitric oxide (NO) or N_2_O by denitrifying organisms and consequently be lost to the atmosphere. Despite the apparent importance of nitrification in the generation of N_2_O and NO_3_^-^, the relative contributions of comammox, AOA, AOB and NOB populations in this process, especially during soil fertilization events, is less clear (6). Advancing this issue is essential for better prediction of the contributions of these microbial taxa to the nitrogen cycle and the modeling of the corresponding activities and products. High-throughput sequencing and proteomic approaches offer the means to characterize the nitrogen pathways in the environment. However, to what extent these omic approaches reflect process rates is still unclear.

Although DNA, RNA, and protein abundances all reflect microbial potential and responses to environmental changes and thus, can be used to study nitrogen cycling in soils, each measurement generally offers different types of information. For instance, metagenomics (DNA level) offers a comprehensive overview of the functional potential of microbial communities but does not generally reflect active community members or functions. Short-term microbial responses to external changes (e.g., nitrogen addition) can be tracked by analyzing the actively expressed genes (i.e., metatranscriptomics). For instance, the relationship between measured nitrification processes and the ammonia monooxygenase (*amoA*) transcripts have revealed differences between archaeal and bacterial activity in acidic soils (7). Proteomics provides a third level of molecular information much closer to the metabolic processes by reflecting synthesized enzymes that catalyze reactions. Although proteomics has been applied to only a limited number of natural microbial communities, the results have provided new insights about metabolic pathways and interdependencies among microbial groups [reviewed in (8)]. Furthermore, recent advances in metagenomics and metaproteomics techniques as well as integration with isotope-based technologies (e.g., NanoSIMS) have disentangled the role of previously elusive keystone microbial populations. The combined application of metagenomics and metaproteomics has provided new understanding of novel not yet cultured microorganisms participating in the cycling of sulfur, nitrogen, and carbon in the terrestrial subsurface (9).

Only a few studies have examined how the above approaches correlate with process rates, especially in soil ecosystems that are characterized by low metabolic activity along with high microbial diversity and heterogeneity. Thus far, almost all studies have provided only qualitative results from applications of omics to soils (10). Quantitative results in a few recent reports have focused mostly on systems with reduced diversity or specific functions and taxa (as opposed to community-wide activities). For instance, metatranscriptomic approaches examining the degradation of the herbicide atrazine by *Escherichia coli* in bioreactors revealed a linear relationship between the measured enzymatic activity and the transcripts encoding the associated enzyme (11). Additionally, in microbial leaf litter decomposition incubations, cellulase and xylanase protein abundances were positively correlated with their corresponding enzymatic activities (12). On the other hand, even though the combination of multi-omic datasets provided new insights into diversity and gene potential of microbial communities of permafrost ecosystems, the datasets were less predictive of measured process rates (13). Therefore, to what extent the omic measurements correlate with each other and with process rates in soils remain unclear.

Toward closing this knowledge gap, we examined nitrogen-amended sandy soils obtained from a site with a history of agricultural management and application of synthetic nitrogen fertilizer. A prior year-round characterization of field samples from the same agricultural site revealed increased abundance of novel *Thaumarchaeota* and comammox nitrifiers, but the findings were limited to metagenomics (14). Here, our goal was to assess the strengths and limitations of multi-omics in detecting microbial activity by correlating measurements of DNA, RNA, and protein abundances with measured rates of nitrate formation and N_2_O production in soils incubated under controlled conditions in the laboratory. The results reveal that metatranscriptomic data best reflected the measured nitrification rates under the tested experimental conditions.

## RESULTS

### Nitrification activity in soil microcosms

We first examined nitrification activity in nitrogen-amended microcosms with an equimolar mixture of NH_4_^+^ and urea during an eight-day period by following NO_3_^-^ formation and NH_4_^+^ disappearance. Based on the NH_4_^+^ concentration patterns, urea quickly hydrolyzed to release NH_4_^+^ within the first two days of incubation (Figure 1a). Specifically, the NH_4_^+^ concentrations peaked at 48 hours of incubation (18.02 ± 1.5 µgN-NH_4_^+^ g^-1^ dry soil) from urea hydrolysis, and decreased to 5.4 ± 2.5 µgN-NH_4_^+^ g^-1^ dry soil by 192 hours of incubation due to nitrification. Nitrification activity increased five to eight days after the addition of the NH_4_^+^ and urea mixture, reaching an average rate of 4.1 ± 0.87 µg N-NO_3_^-^ g^-1^ dry soil d^-1^ (n = 6) after 192 hours of the incubation (Figure 1b). The NO_3_^-^ concentrations gradually increased from an initial value of 0.81 ±0.28 µgN-NO_3_ g^-1^ dry soil to 1.91 ± 0.5 at 120 hours of incubation, and then increased at a faster rate to 15.06 ± 2.7 µgN-NO_3_^-^ g^-1^ dry soil at 192 hours of incubation (Figure 1a). As a result of nitrification activity, pH values decreased across replicated nitrogen-amended microcosms during the incubation (Supplementary Table 1). In order to examine the generation of N_2_O possibly generated as a by-product of oxidation reactions during nitrification, we measured the production of N_2_O in nitrogen-amended incubations. Net N_2_O production rates in the incubation headspace increased from 0.08 ± 0.006 ng N-N_2_O g^-1^ dry soil d^-1^ after 24 hours to 0.71 ± 0.57 ng N-N_2_O g^-1^ dry soil h^-1^ at the end of the incubations (Figure 1c). Control microcosms receiving only irrigation water (i.e., no nitrogen amendment) did not show net NH_4_^+^oxidation.

**Figure 1.**
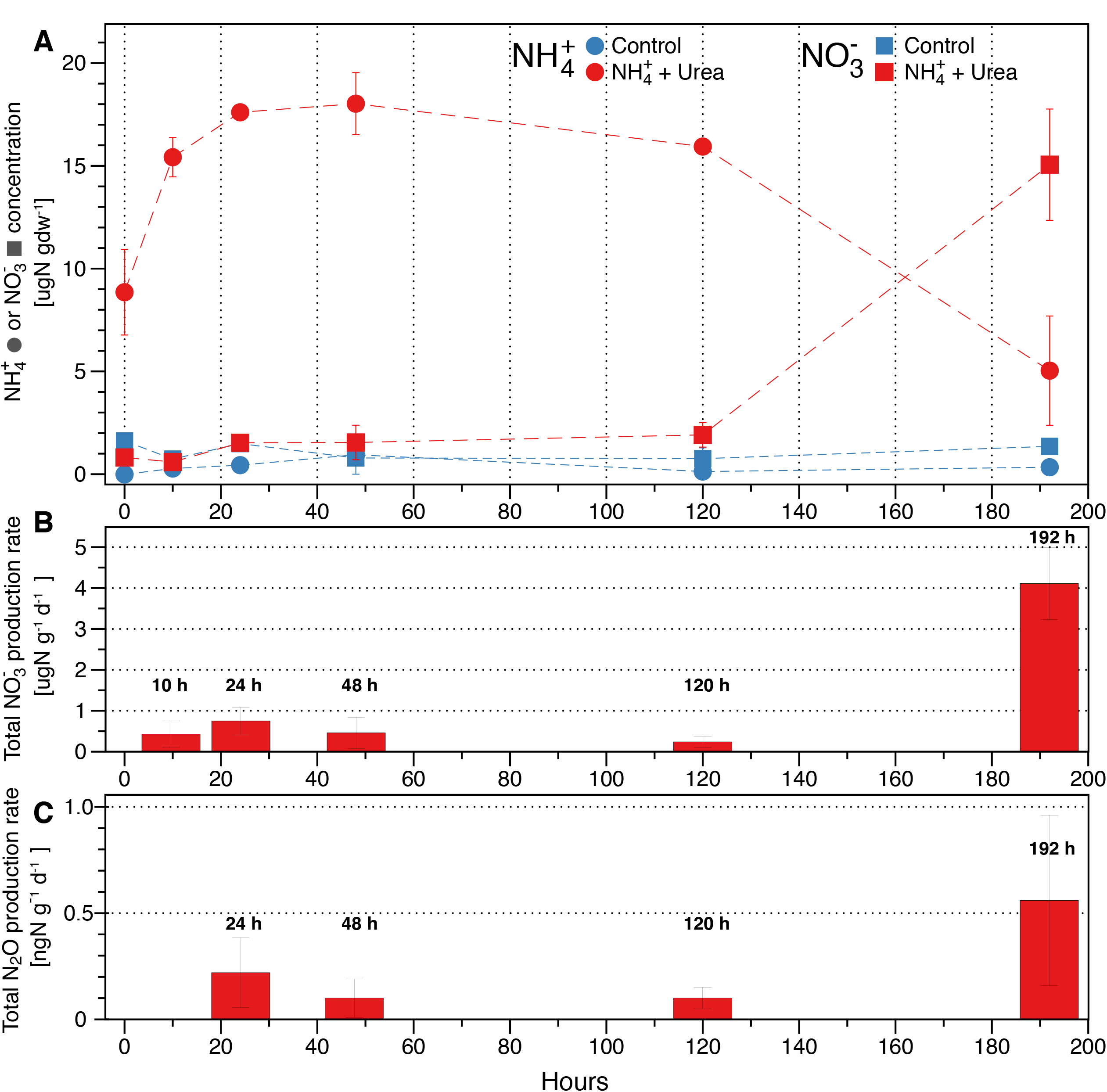
Nitrogen pools and fluxes in soil incubations amended with NH_4_^+^ and urea. Mean NH_4_^+^ and NO_3_^-^ concentrations (A), total NO_3_^-^ production rate (B), and total N_2_O production rate (C) for the nitrogen-amended and control (irrigation water only) microcosms at each incubation time point. Error bars represent the standard deviation from replicate samples (n=6 for nitrogen-amended and n=3 for control).

To evaluate possible differences between the use of NH_4_^+^ or urea in the nitrifying activity, we determined ^15^NO_3_^-^ production rates using nitrogen stable isotopes in the microcosms. In general, ^15^NO_3_^-^ production was similar between 15NH_4_^+^ and 15N-urea microcosms, although rates were higher after 10 and 48 hours of incubation (two tailed *t*-test, *P*<0.01) in ^15^NH_4_^+^ and 15N-urea microcosms, respectively, but converged thereafter (Supplementary Figure 1). By the end of the incubations, approximately half of the added ^15^N was converted to ^15^N-NO_3_^-^ (49-55% for both labelled solutions/treatments), and only a small fraction converted to ^15^N-N_2_O (0.006-0.01%). The remaining added nitrogen was presumably converted to N_2_, assimilated into microbial biomass, or adsorbed to soil particles.

### Soil metagenomes and metatranscriptomes

To explore the genetic potential of microbial communities in control and nitrogen-amended microcosms, we examined the metagenomes and metatranscriptomes obtained from the incubated soils. Metagenomes ranged from 23.7 to 53.4 and metatranscriptomes from 10.1 to 31.3 million short-reads per sample (Supplementary Tables 2 and 3). The estimated average coverage based on read redundancy using Nonpareil (15) ranged from 0.27 to 0.42 for the soil metagenomes (values range from 0 to 1). The co-assembly of selected soil metagenomes generated 1.52 million contigs over 500 bp (assembly N50=1,176) and 1.56 million predicted protein-coding genes.

A high fraction of ribosomal RNA was detected for all metatranscriptomes ranging from 94% to 98% of the total sequences (Supplementary Table 4). No rRNA depletion step was performed during our metatranscriptomic protocol due to overall low total RNA yields from the soils. As expected based on the length of the rRNA genes, 23S rRNA/16S rRNA ratios ranged from 1.7 to 1.9, indicating adequate RNA quality. Bacterial 16S rRNA (16S) was the most abundant, ranging between 30.6% and 35.9% of total transcripts per sample. Archaeal 16S and eukaryotic 18S rRNA molecules were less abundant, with values ranging from 0.09% to 0.15% and 0.55 to 2.9%, respectively.

### Taxonomy of microbial soil populations based on 16S rRNA gene sequences

The taxonomic composition and abundances of the main microbial groups determined from recovered 16S rRNA (16S) gene sequences (DNA level) from nitrogen-amended incubations, were generally stable during incubations. At the class taxonomic level, *Actinobacteria*, *Betaproteobacteria*, and *Gammaproteobacteria* were the most abundant groups in metagenomes, accounting for more than 57% of the total community in nitrogen-amended incubations (Supplementary Figure 2). The taxonomic composition derived from metatranscriptomes (cDNA reads) was also stable during the incubations but the abundances for main taxonomical groups were substantially different from the metagenomes. For instance, *Betaproteobacteria*, *Gammaproteobacteria*, and *Flavobacteria* were among the most abundant groups in cDNA samples, accounting for an average of 77.5% of the 16S transcripts. In agreement with our previous results based on field samples from the same agricultural site (14), bacterial and archaeal groups associated with the nitrification processes were comparatively less abundant than the aforementioned groups in both DNA and cDNA datasets. For instance, known AOB and NOB genera such as *Nitrosomonas* and *Nitrospira* had average relative abundances of 0.01% and 1.6% of the total populations in the metagenomes from incubated soils. Additionally, the relative abundances of the AOA genera related to *Nitrososphaera* and *Nitrosopumilus* were 0.9% and 0.3% in the microcosm metagenomes. Similar low abundances were determined for known nitrifier genera in metatranscriptomes. Notably, 16S transcript abundances for AOB conspicuously increased during the incubation period (Supplementary Figure 3). In fact, relative 16S gene expression ratios (cDNA/DNA) for AOB and NOB belonging to *Nitrosospira*, *Nitrosomonas* and *Nitrospira* increased 3-,6-, and 14-fold between 10 and 192 hours. In contrast, the 16S gene expression levels for the archaeal groups *Nitrososphaera* and *Nitrosopumilus* were stable during the same incubation period, although with a slight increase in relative expression at 48h of incubation (Supplementary Figure 3).

### Individual populations from microcosm metagenomes

The assembly and binning of the soil metagenomes recovered 11 metagenome-assembled genomes (MAGs) mostly representing *Proteobacteria*, *Acidobacteria*, *Actinobacteria* and *Nitrospirae* phyla. Most of the recovered MAGs represented novel genera (n=7) and species (n=5) when the taxonomic novelty was evaluated against 10,487 reference genomes (taxonomically classified at the species level) using genome-aggregate amino acid identity (AAI) thresholds for taxonomic rank delineation (16) (Supplementary Table 5). Given that none of the MAGs represented AOA, AOB, NOB, or comammox populations, we included MAGs obtained from a previous analysis of field samples from the same site (Havana county, Illinois, USA) and depth as the soil used in the soil microcosms in the present study (14). MAGs potentially involved in nitrification processes were likely missed in the microcosm metagenomes due to comparatively lower sequencing effort or because of sample heterogeneity (e.g., lower population abundance) but had relatively higher abundance in the previous field samples. The MAGs (designated with the letter F at the end of their name for Field metagenomes) consisted of two complete ammonia oxidizer (comammox) *Nitrospira* MAGs (MAG021F and MAG017F) and five ammonia-oxidizing archaea MAGs representing the *Thaumarchaeota* lineages I.1b (MAG032F and MAG019F) and I.1a (MAG004F, MAG109F, and MAG001F) (Supplementary Figure 4b). The *Nitrospira* MAG007, obtained from the microcosm metagenomes, was closely related to previously described soil comammox (e.g., MAG017F) organisms, sharing 67.8% AAI (SD: 18% based on 2201 shared proteins). However, the MAG007 only encoded a hydroxylamine oxidoreductase (*haoA*) gene and lacked *amoA* and *nxrA* genes (Supplementary Figure 4b). Furthermore, the *Nitrospira* MAG007 formed an independent but related cluster to the soil comammox organisms when reconstructed phylogenies using concatenated single-copy genes were evaluated (Supplementary Figure 5). Thus, AAI values and phylogenetic reconstruction supported the affiliation of MAG007 to *Nitrospira*, but the lack of genes involved in ammonia and nitrite oxidation (possibly due to low sequencing coverage) made it inconclusive whether this taxon is involved in nitrification processes and might indicate divergence from previously described soil comammox organisms.

Relative expression values of MAGs (measured as transcripts or reads per kilobase million, RPKM) were used as a proxy for comparing the response and metabolic activity among nitrifying bacteria and archaea during incubations. Even though expression values for most nitrifying MAGs belonging to *Nitrospira* and *Thaumarchaeota* were stable and relatively low, AOA MAGs 004F, 019F and comammox MAG017F, had, on average, the highest expression values throughout the incubations (Supplementary Figure 4a). For instance, the increase in expression values for AOA MAGs belonging to the I.1b clade, 004F and 032F, were 39% and 50% after 48 hours of incubation (compared to expression levels at 10 hours incubation), respectively. In contrast, gene expression of comammox MAG017F and *Nitrospira* MAG007 increased by 59% and 68% after 120 and 192 hours of incubation, respectively (Supplementary Figure 4a). Note that AOB and NOB were not included in the RPKM analysis due to lack of recovered MAGs representing these populations (see above). Nonetheless, a gene-based approach allowed the analysis of changes in transcript abundances of genes involved in nitrification activity for AOB and NOB nitrifiers (see below).

### Quantification of nitrification genes in microcosms

To further explore the microbial nitrification processes in incubated soils at the gene level, we specifically quantified gene fragments and transcripts directly involved in nitrification reactions. Relative expression values belonging to the gene encoding urease subunit c (*ureC*) were stable throughout the incubation but average abundances were relatively low compared to other nitrification genes (Figure 2a). The relative expression of the bacterial gene encoding ammonia monooxygenase subunit alpha (*amoA*) was 53.5-fold higher compared to the expression values at 10h of incubation. Most of the detected *amoA* transcripts (cDNA) were phylogenetically affiliated with *Betaproteobacteria* and corresponded to up to 90% of the total detected bacterial *amoA* transcripts at 192 hours of incubation (Figure 2a). Unlike betaproteobacterial *amoA*, abundance of transcripts belonging to comammox were stable throughout the incubation. Furthermore, transcripts belonging to comammox were more abundant compared to *Betaproteobacteria amoA* transcripts after 48 hours of incubation; however, at 192 hours of incubation, betaproteobacterial *amoA* DNA abundance increased 66-fold, whereas comammox *amoA* gene fragments remained stable (Figure 2b). The latter results indicated that the comammox *amoA* may be more abundant under field conditions but betaproteobacterial *amoA* might show a faster response upon ammonia addition, which was also consistent with a previous study (14). Although the relative expression for the archaeal *amoA* was more stable throughout the incubation compared to its betaproteobacterial counterparts, a maximum expression was reached after 120 hours of incubation, suggesting that archaeal AmoA activity temporarily increased at later time points during the incubation. Archaeal *amoA* transcripts belonging to the group I.1b were ~7 times more abundant than their I.1a counterpart across the incubations (Figure 2b). Similar to *amoA* patterns, the relative expression for the betaproteobacterial hydroxylamine oxidoreductase (*haoA;* NH_2_OH→NO_2_^-^) steadily increased during the incubations, whereas comammox *haoA* transcripts represented the remaining smaller transcript fraction and were stable throughout the incubations (Figure 2b). Expression values for the nitrite oxidoreductase subunit alpha (*nxrA;* NO_2_^-^→NO_3_^-^) had a 12.4-fold increase compared to the 10-hour time point, consistent with the patterns observed for the previous nitrification genes and NO_3_^-^ accumulation. Unexpectedly, expression values for *nirK* (NO_2_^-^→NO) affiliated to *Thaumarchaeota* were higher compared to *nirK* transcripts assigned to the *Nitrospira* clade. In fact, *Thaumarchaeota nirK* transcripts had a 3.4-fold increase after 192 hours of incubation relative to earlier sampling points, indicating that *Thaumarchaeota* might have been more active in the reduction of nitrite compared to other steps of nitrification. Specifically, there was a 3.4-fold increase for clade I.1b *nirK* transcripts during the 10 to 192 hours of incubation period, whereas the abundance of transcripts from clade I.1a were stable throughout the incubations (Figure 2b).

**Figure 2.**
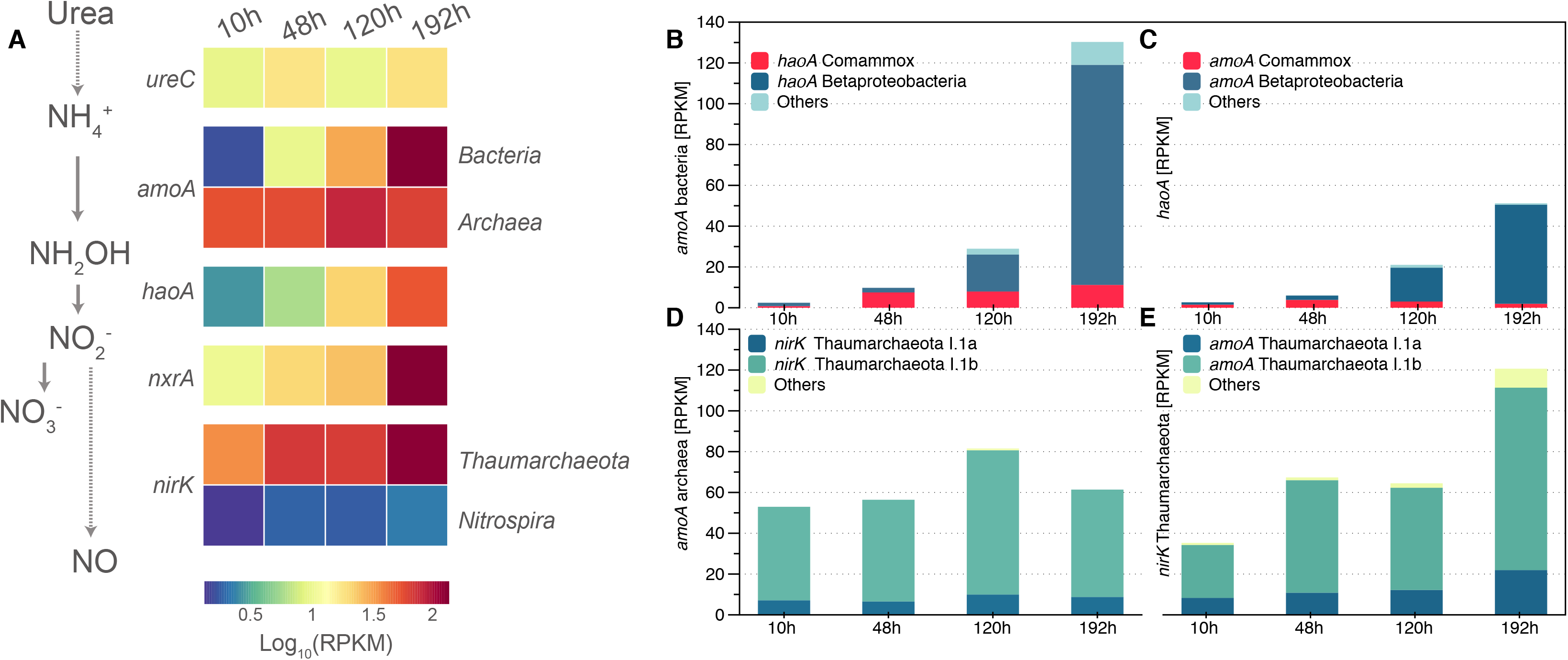
Nitrification genes in incubated soils. A. Relative expression ratios for each nitrification step in incubated soils were determined at 10, 48, 120, and 192 hours incubation. B and C show determined RPKM values for bacterial *amoA* (B), *hao* (C), thaumarchaeotal *amoA* (D), and *nirK* (E) transcripts from metatranscriptomes.

In summary, the metatranscriptomic profiles suggested that AOB, but not comammox, responded rapidly to the nitrogen amendment, whereas AOA followed with less pronounced transcriptome shifts. The response of AOB, and to a lesser extent AOA, was also reflected at the DNA level, albeit with a substantial time delay. For instance, shifts were observed early at the transcript level while at the DNA level, changes were mostly evident 192 hours after the start of incubation (Supplementary Figure 6a, b). These results were consistent across the individual nitrification steps and indicated that at least the AOB nitrifiers grew in response to nitrogen addition.

### A proteomic perspective in soil microcosms

A metaproteomic analysis of the control and nitrogen-amended microcosms at 192 hours of incubation detected a total of 2,892 and 1,629 non-redundant peptides, respectively. A total of 844 peptides were shared among control and nitrogen-amended incubations, whereas 2,048 and 785 were exclusively present in each microcosm, respectively. Most of peptides detected in control and nitrogen-amended incubations matched protein sequences predicted from metagenomic assemblies (89.4% and 88.2%, respectively) and the remaining fraction matched reference proteomes (Supplementary Table 6). The top 20 most abundant proteins in control and nitrogen-amended treatment microcosms were related to housekeeping and transport proteins whereas in the latter incubation, oxidoreductases for small carbon and alcohol molecules and ATP synthesis were among the most abundant proteins detected (Supplementary Table 7). The taxonomic affiliation, at the class level, for the most abundant annotated peptides belonged to *Alphaproteobacteria*, *Betaprotebacteria*, and *Acidobacteria* in control and nitrogen-amended incubations. Although there were major compositional changes for abundant groups such as *Betaproteobacteria* (40% decrease) and *Gammaproteobacteria* (50% decrease) (Figure 3a), increased abundance was detected for less abundant groups commonly associated with the nitrification process. For instance, close to a 2.2-fold increased abundance for nitrogen-amended incubations were detected for peptides belonging to *Nitrospira*. Detected peptides related to folding and synthesis were the most abundant and had similar abundances in the control and nitrogen-amended microcosms after 192 hours of incubation. However, the relative abundance of ATP synthases and transcription categories were higher in the nitrogen-amended samples relative to the control, presumably as a consequence of a higher microbial activity generated after the nitrogen input. On the other hand, heat-shock and degradation proteins were more abundant in the control incubation, probably reflecting a more prevailing dormant state for the microbial communities in these samples (Figure 3b). However, unlike the metagenomic and metatranscriptomic datasets, only some peptides involved in nitrification were identified using metaproteomics. For instance, the detected peptides directly involved in nitrification pathways corresponded to the nitrite oxidoreductase subunit B (NxrB), which had a 31.3% abundance increase in the nitrogen-amended samples compared to the control.

**Figure 3.**
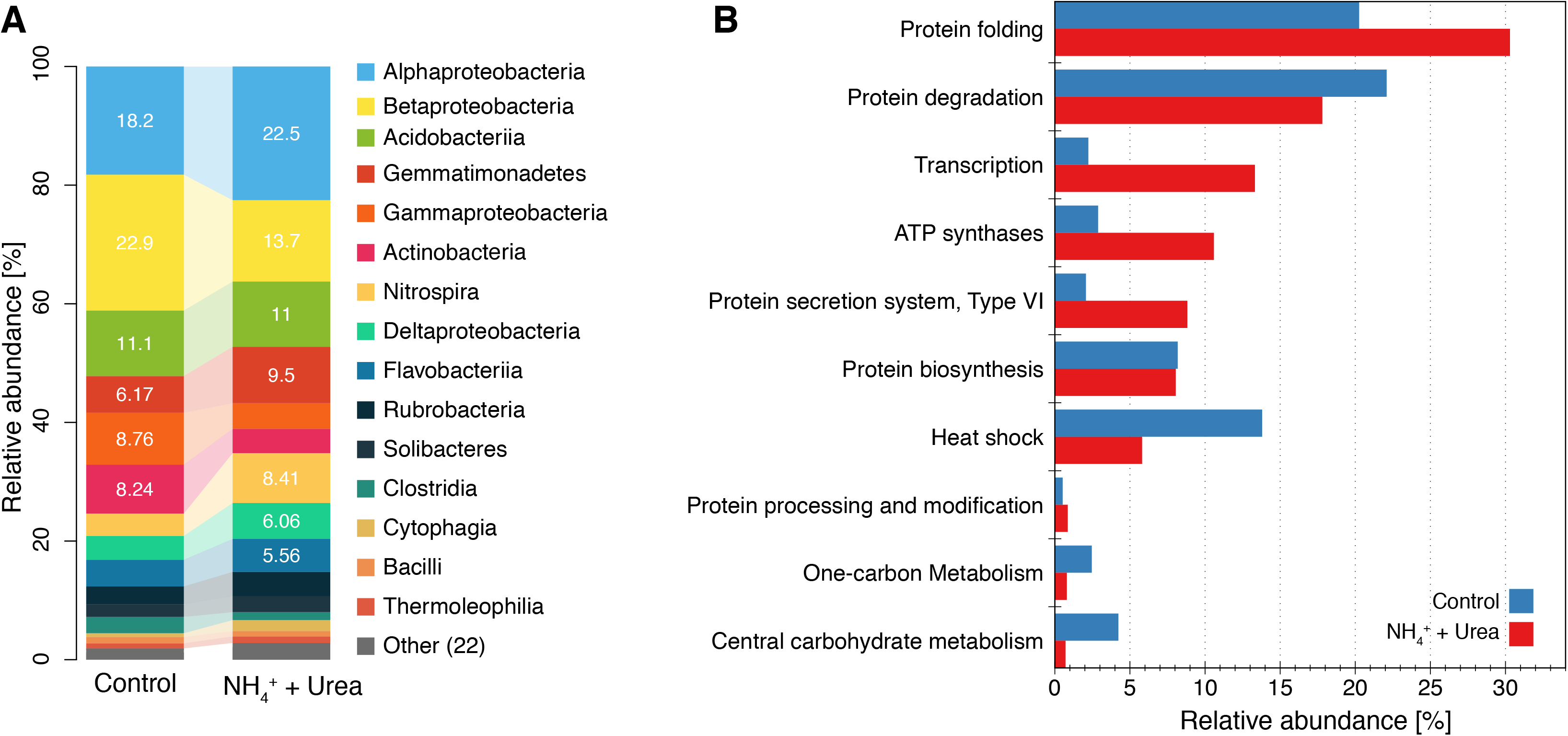
Metaproteomic analyses of incubated soils at 192 hours of incubation. Panel A shows taxonomic affiliation (class) and abundance (average spectral counts) for peptides detected in control and N-amended incubations. Panel B shows summarized functional annotation of detected peptides using SEED functional categories.

## DISCUSSION

### Using multi-omic approaches for examining process rates

Measuring nitrification rates in incubated soils allowed us to evaluate the explanatory and predictive power of omic approaches in a highly diverse soil system. Despite all three omic approaches revealed increased abundance for target genes, transcripts and proteins related to nitrification pathways, they differed in temporal resolution and quantitative capabilities. For instance, the strongest agreement to the observed nitrification processes (i.e., ammonia or nitrite oxidation) was for the metatranscriptomic data within the first days of incubations (e.g., Figure 2b), whereas metagenomes lagged behind and only reflected the ongoing nitrification process after 192 hours of incubation (e.g., Supplementary Figure 6a, b). These data were presumably attributed to the fact that growth (e.g., at least a few replication cycles) should occur before metagenomics can reveal shifts in relative abundance over time. Note that microbial growth was not explicitly measured by our study to further corroborate the above conclusions and interpretations. Therefore, metagenomics could also reflect underlying microbial processes if the processes are ongoing for a period of time and are coupled with the growth of the corresponding organisms. In contrast, if the goal is to see immediate responses to a perturbation or the perturbation is short-lived (e.g., lasting a few hours), metatranscriptomic data will be preferable. We also observed that metatranscriptomes were as good as metagenomics, if not better, at reflecting microbial activity for nitrification processes even at later incubation time points. In contrast, the metaproteomes offered, at most, a qualitative glimpse at nitrification processes and were less definitive in identifying common nitrification markers. The latter was largely attributable to the computational challenges associated with proteomic data such as high peptide redundancy and the requirement of high-quality assemblies which are still challenging for highly complex soil metagenomes. Furthermore, many challenges remain for efficient extraction of membrane proteins from low abundance organisms such as nitrifiers. Ultimately, these technical limitations could be reflected in a lower number of detected proteins compared to the number of metagenomic and metatranscriptomic reads recovered that encoded the proteins of interest.

While shifts in 16S gene ratios (cDNA/DNA) were relatively small for AOA, the 16S and functional gene ratio shifts (e.g., *amoA*) for AOB/NOB were much more pronounced throughout the incubations (Supplementary Figure 3). Nonetheless, there were changes in transcript abundances for nitrification genes from both microbial groups in the microcosms (Figure 2). These results might reflect an active and growing state for AOB/NOB and mostly active AOA communities as observed before for agricultural soil microcosms (17). The differences observed between target gene abundances and 16S gene ratios from AOA could reflect a limitation of the latter data when used as a proxy for assessing microbial activity (18). However, more frequent sampling and incubations under different physicochemical conditions will be required for more robust conclusions to emerge on the exact relationship(s) between molecular level information and process rates. The results reported here provided an overview of this relationship for soils and are highly promising for the future.

In terms of the ecological adaptation of the nitrifiers analyzed here, the Havana agricultural site has had a long history of cyclical seasonal inputs (e.g., fertilizers) that have shaped the structure of microbial communities in different soil layers. The AOA and AOB communities in the Havana site have legacy establishments at the 20-30 cm soil depth and are under relatively stable environmental conditions compared to the top soil layer (14). Thus, nitrogen amendments tested in our experiment and experimental conditions might not represent closely the conditions usually experienced by the examined AOA and AOB communities. The rapid response of AOB observed here might be a reflection of physiological adaptations of AOB to thrive under high nitrogen content as reported previously (17). In contrast, the low response observed for comammox and some AOA communities might reflect their limited physiological capabilities to respond to high nitrogen concentrations (2, 3) that were assayed in our experimental setup.

Previous authors have also found metatranscriptomic approaches to be better predictors of measured microbial activity (11) in controlled laboratory systems amended with exogenous organic compounds, but have been more limited in providing insights into the whole-microbial community response to the amendment. For instance, the changes in transcripts observed at early incubation points for specific lineages (e.g., comammox vs. betaproteobacterial *amoA*) suggested ongoing microbial activity that became evident only at the DNA level (relative abundance) at the last incubation point in the metagenomes (Figure 2b). Future incubation studies could shed light on the intrinsic differences between nitrifier (and denitrifier) communities by testing variables such as oxygen availability (i.e., water saturation) and different agricultural soil types. For instance, the incubation conditions used in our study deliberately promoted nitrification over denitrification processes and as a result, the N_2_O production was detected due to the former process. Consequently, nitric oxide (e.g., *norB*) and nitrous oxide reductases (e.g., *nosZ*) transcripts, which are responsible for N_2_O production and consumption during denitrification, respectively, were not detected in our metatranscriptomes datasets (i.e., abundance below detection limit). Also, the use of nitrification inhibitors could help to elucidate the origin of the measured N_2_O whether production was biotic or abiotic, for which our data are limited in predicting. Thus, the integration of *in situ* rates along with the microbial dynamics examined by metatranscriptomes and metagenomes could provide the means to better understand and predict nitrification and N_2_O emission in agricultural soils.

### New insights into nitrification pathways

The metagenomic and metatranscriptomic datasets combined with phylogenetic approaches provided a closer examination of the poorly studied microbial diversity in agricultural soils. Assessing the individual gene level, as opposed to whole genome transcript level, provided more robust results for relating population response to measured nitrification reactions, presumably due to higher sequence coverage (less noise). Even though a direct comparison at the genome level between AOB, NOB and comammox AOB was not possible due to the lack of recovered MAGs representing AOB and NOB populations, analysis of individual 16S and nitrogen cycling genes elucidated the importance of AOB and NOB in the microcosm experiments. Our results showed that even though betaproteobacterial *amoA* transcripts responded to the addition of ammonium and urea, the relative abundance of comammox *amoA* transcripts was stable (i.e., not responding to the nitrogen amendment), although comammox populations were relatively more abundant than AOB in the microcosms. This observation is consistent with previous metagenomic results from the same agricultural soil, where comammox *amoA* genes and the organisms encoding these genes represented the highest fraction of nitrifying bacteria (14). The differences between measured genes and transcripts indicated that the incubation conditions favored the activity of *Betaproteobacteria* over comammox nitrifying bacteria, suggesting ecophysiological differences among these taxa for the incubation conditions or added substrates compared to field conditions.

The sequencing of isolates and environmental AOA genomes has shown that even though they encode an AmoA protein, they lack a canonical hydroxylamine oxidation pathway (19). Previous studies have proposed that nitric oxide is essential for hydroxylamine oxidation to nitrite in archaea (20). The proposed mechanism involves oxidation of ammonium to hydroxylamine followed by oxidation to nitrite catalyzed by a putative Cu-protein that uses nitric oxide as co-reactant for the oxidation of hydroxylamine. Interestingly, nitric oxide has been proposed to be derived from the activity of the NirK enzyme present in all AOA sequenced genomes. Our results show that unlike AOA *amoA* or bacterial *nirK* transcripts, *Thaumarchaeota nirK* transcripts increased in abundance in the incubated soils, supporting the abovementioned hypothesis. Therefore, even though AOA *amoA* transcripts did not show clear changes in abundances compared to their bacterial counterparts, these results might be in agreement with the previous hypothesis, and likely denote an unaccounted role for *Thaumarchaeota nirK* in nitrification in agricultural soils.

### Multi-omic limitations

Soil samples are challenging to analyze not only because of their heterogeneous structure and chemical composition, but also because of the highly diverse microbial communities and slow growth kinetics. Despite the advancements presented here, there are still opportunities for further improvements. For instance, here we analyzed total RNA extractions from soils where ribosomal rRNA transcripts represented 94-98% of the total sample, limiting our study to a small fraction of transcripts related to functional genes. Current experimental approaches offer successful rRNA depletion for environmental samples, when RNA yields are not limiting (21). Additionally, all the results represented here provide only relative abundances for measured microbial markers. For instance, approaches such as qPCR or internal standards spiked into the DNA or cDNA library for sequencing (21) can strengthen and provide improved quantification compared to those presented here.

Metaproteomics offered an additional layer of information for the microbial activity, but it was less comprehensive compared to metagenomes and metatranscriptomes. Even though our database for proteomic analyses included a high fraction of nitrification proteins predicted from these agricultural soils, only peptides belonging to the NxrB were detected. The results obtained were attributable, at least partially, to the low biomass, especially for the low abundance nitrifiers targeted here. Further, possible protein extraction biases due to the complexity of soil matrices as well as limited extraction of membrane proteins, such AmoA, might have also influenced the outcome of our efforts (22). Nonetheless, the abundances for several peptides belonging to housekeeping proteins of nitrifier organisms were increased during the incubation time, consistent with the results from metagenomic and metatranscriptomic approaches. Therefore, metaproteomics provided a qualitative confirmation of the underlying nitrification processes ongoing during our incubations and of the responsible taxa. Alternative proteomic approaches focused on a preselected set of proteins (i.e., selected reaction monitoring or target proteomics) could be used to explore low abundance nitrification proteins. For instance, targeted proteomic approaches have been used to study proteins in low abundance involved in bioremediation pathways in highly-diverse environmental systems (23). Therefore, targeted proteomics might offer new opportunities for researchers interested in detecting low-abundance peptides and prediction of process rates in complex samples (24).

The analyses of different omic levels obtained from the incubations showed a high correspondence between nitrification gene markers and nitrification process rates. The gene fragments and transcripts were mostly affiliated to novel nitrifier populations similar to those previously described in field soil metagenomes from the same agricultural site (14). Therefore, the gene and genome sequences reported here could facilitate future investigations of nitrogen cycling in agricultural fields; for instance, by applying qPCR assay targeting the key taxa and biomarker genes and transcripts. The combination of metagenomic and metatranscriptomic approaches used in our study provided a promising strategy for examining microbial activity in agricultural soil environments. Therefore, the findings presented here highlighted the potential of omics data to serve as reliable proxies for examining microbial processes *in situ*, especially in soils, which has been proven to be among the most challenging tasks for environmental studies.

## MATERIALS AND METHODS

### Soil Sampling

Our study was focused on an agricultural plot located in the Havana County, Illinois, USA (lat 40.296, long 89.944; elevation, 150 m). The site is representative of the US Midwest and has a long history of conventionally managed corn and soybean crop rotation. In October 2014, we collected ~2 kg of bulk soil from a 20-30 cm soil depth as previous results have shown significant presence of ammonia-oxidizing microorganisms in this layer (14).

### Soil Incubations, Gas and Chemical Analyses

Soil microcosms were established in triplicates, using ~120 g of soil (~8% moisture content) in 500 ml gas-tight canning jars equipped with gas sampling ports, and were sampled at six time points (0, 10, 24, 48, 120, and 192 hours). To set up the microcosms, 6 ml of 40 mM NH_4_Cl and 20 mM urea (80 mM N) in water used for irrigation at the site was added to two separate batches of 400 g of soil (Final concentration= 1.2 µmoles-N/g or 18.3 µg-N/g dry weight). Two stable isotope treatments were done, one for NH_4_Cl (50% ^15^N-NH_4_Cl and 50% ^14^NNH_4_Cl) and one for urea (50% ^15^N-NH_2_CONH_2_ and 50% ^14^N-NH_2_CONH_2_). The two treatments allowed for differentiating how the products of nitrification differed between urea and NH_4_Cl when both were present. After vigorously mixing, 120 g were dispensed into three separate microcosm jars and incubated in a dark growth chamber with diurnal temperature fluctuation of 22-24 °C as observed in Havana field soil at 20-30 cm during the spring fertilization period (early June). Triplicate microcosms each receiving 6 ml of filtered irrigation water (no nitrogen amendment) served as controls. After each sampling point, headspace gas was collected from closed jars and the N_2_O concentration was measured on a Shimadzu GC-2014 gas chromatograph (Columbia, MD) equipped with an electron capture detector. Jars were opened for soil sampling and to reestablish equilibration with atmospheric air before being resealed until the next sampling. Residual ammonium and nitrate in soil subsamples (20 g) were extracted in 2 M KCL and the concentrations were determined using colorimetric analysis on a flow injection auto-analyzer (Lachat Instruments, Milwaukee, WI) (25). Soil pH (1:1 in water) and gravimetric water content were measured at each time point (Supplementary Table 1). ^15^N isotopic composition of N_2_O in collected jar headspace samples was determined using an IsoPrime 100 isotope ratio mass spectrometer interfaced with an IsoPrime trace gas analyzer (Cheadle Hulme, UK) at the University of Illinois at Urbana-Champaign. The ^15^N atom % enrichment of the NO_3_^-^ pool was determined using acid trap diffusion (26) and analysis of the diffusiondisks on a Vario Micro Cube elemental analyzer (Elementar, Hanau, Germany) interfaced to an IsoPrime 100 continuous flow isotope ratio mass spectrometer (Cheadle Hulme, UK). 15NO_3-_and ^15^N_2_O production rates were calculated from the change in ^15^NO_3_^-^ and ^15^N_2_O concentrations, respectively, from one time point to the following sampling time point. NO_3_^-^ and N_2_O production rates were estimated from the ^15^NO_3_^-^ and ^15^N_2_O production rates based on the mean ^15^N excess atom % of the NH_4_^+^ source pool (27). No inhibitors of nitrogen cycle pathways were used in the incubations.

### Nucleic Acid Extractions

DNA was extracted from ~0.5 g of soil using a modified phenol-chloroform and purification protocol as previously described (28). For RNA extraction, 2 gr of soil was preserved in LifeGuard (MoBio) and stored at −80°C. A modified protocol derived from the PowerMax Soil DNA kit for extracting RNA was used for total RNA extractions (MoBio). TURBO DNAse (Ambion) was used to remove DNA according to the recommendations of the manufacturer. Nucleic acid extracts were quantified using Quant-it ds DNA HS and HS RNA assays (Invitrogen) according to the instructions of the manufacturer. RNA quality was assessed using Agilent RNA 6000 pico kit (Agilent Technologies) and samples having RNA integrity number (RIN) above 7 were used.

### Nucleic Acid Sequencing

For metagenomes, dual-indexed DNA sequencing libraries were prepared using the Illumina Nextera XT DNA library prep kit according to manufacturer’s instructions, except that the protocol was terminated after isolation of cleaned amplified double stranded libraries. For metatranscriptomes, single-indexed cDNA sequencing libraries were prepared using ScriptSeq v2 protocol using ~25 ng of total RNA as input. All DNA and cDNA library concentrations were determined by fluorescent quantification using a Qubit HS DNA kit and Qubit 2.0 fluorometer (ThermoFisher Scientific) according to manufacturer’s instructions and samples were run on a High Sensitivity DNA chip using the Bioanalyzer 2100 instrument (Agilent) to determine quality and average library insert sizes. An equimolar mixture of the libraries was sequenced on an Illumina HiSEQ 2500 instrument (School of Biological Sciences, Georgia Institute of Technology) for a rapid run of 300 cycles (2 x 150 bp paired end) using the HiSeq Rapid PE Cluster Kit v2 and HiSeq Rapid SBS Kit v2 (Illumina). Adapter trimming and demultiplexing of sequenced samples was carried out by the Illumina software, according to the recommendations of the manufacturer.

### Short-read Analyses

Metagenomic and metatranscriptomic raw reads (FASTQ) for all samples were trimmed using SolexaQA (29) using a Phred score cutoff of 20 and minimum fragment length of 50 bp. Short-reads derived from metatranscriptomes were merged using PEAR using default parameters (30). Average coverage for each sequenced metagenome was determined by Nonpareil (15) using default settings except that 2,000 reads were used as query (-X option) (Supplementary Tables 3 and 4).

Short-read sequences encoding 16S rRNA gene fragments were extracted from each metagenome and metatranscriptome by SortMeRNA (31) and their taxonomy was assigned using RDP classifier (cutoff 50) (32).

To identify and quantify reads encoding specific protein sequences of interest, we used the previously published protein sequences as references (14) for the archaeal ammonia monooxygenase alpha subunit (AmoA), bacterial AmoA, hydroxylamine oxidase (HaoA), nitrite oxidoreductase alpha subunit (NxrA), nitrite reductase (NirK), nitric oxide reductase beta subunit (NorB), nitrous oxide (NosZ), nitrite reductase (NrfA) and DNA-directed RNA polymerase subunit beta (RpoB). Independent ROCker (33) models (length=125 bp) were subsequently built based on these reference protein sequences with the exception of NarG and NxrA, where the sequences were combined into a single model. Trimmed short-reads from soil metagenomes were used as query for BLASTx searches (e-value 0.01) against the latter protein databases and outputs were filtered using the previously generated ROCker models. For metagenomes, target gene abundance in metagenomes was determined as genome equivalents by calculating the ratio between normalized target reads (number of reads matching divided by median protein length) and normalized RpoB reads (number of reads matching divided by median RpoB protein length), a universal single-copy gene. For metatranscriptomes, target transcripts abundance was calculated as reads per kilobase of transcript per million mapped reads (RPKM). Protein databases and ROCker models are available through http://enve-omics.ce.gatech.edu/.

### Assembly and Binning of Metagenomic Populations

Short-read metagenomes from control and treatments (t=0,120 and 192 hours) were co-assembled using IDBA_UD v1.1.1 (34) and binning was performed as previously described (14). Taxonomic classification and degree of novelty (novel species, genus, etc) of the MAGs were obtained from the Microbial Genomes Atlas (MiGA) webserver (35). MAG abundance was determined as the total length of all matching metagenomic or metatranscriptomic reads to the binned contigs from BLASTn searches (identity >=98% and fraction of read aligned >= 50%) divided by the metagenomic or metatranscriptomic sample sizes (in millions of reads) and the length of the bin genomes in Kbp (Kilo base pairs). Reads encoding rRNA sequences (such as 5S, 5.8S, 16S, and 23S) were identified by SortMeRNA, and removed for non-rRNA analyses in order to avoid overestimating abundances.

Phylogenetic reconstruction of MAGs was performed based on the concatenated alignment of universal single-copy proteins identified for each bin using the “HMM.essential.rb” script of the enveomics collection (36). For this, thirty bacterial proteins present in the corresponding bins MAGs were extracted and multiple alignments for each protein were generated using ClustalΩ. Concatenated alignments without invariable sites were generated for archaeal and bacterial alignments using the script “Aln.cat.rb”. Phylogenetic reconstructions were determined using in RAxML v8.0.19 (-f a, -m PROTGAMMAAUTO, –N 100) and visualized in iTol.

N cycle protein sequences in the co-assembly and MAGs were detected using hidden Markov models obtained from FUNGENE (37), using HMMer (38). Detected target N cycle proteins were manually curated, when necessary, by assessing the presence of characteristic amino acid and phylogenetic congruency.

### Phylogenetic Trees and Placement of Short-reads

To assess the phylogenetic affiliation of metagenomic or metatranscriptomic reads, reference and fully assembled protein sequences were aligned using ClustalΩ (39) with default parameters. Resulting alignments were used to build phylogenetic trees in RAxML v8.0.19 (40). Short*-*reads encoding the protein of interest were extracted from metagenomes or metatranscriptomes using ROCker (BLASTx) and placed in their corresponding phylogenetic tree using the methodology previously described (14). Quantification of the number of reads assigned to a specific clade (e.g., to distinguish between *nxrA* or *narG* reads) was done using the “JPlace.distances.rb” script, also available in the enveomics collection. To quantify *nirK* gene fragments assigned to specific clades, the same process as described above was repeated except that all reads detected by multiple ROCker models to previously described clades (41) (clades I+II, III and *Thaumarchaeotea*) were used.

### Shotgun Metaproteomics

Approximately 10 g of soil were collected from the 192 hours control and ^15^NNH_4_^+^ amended microcosms and stored at ^-^80°C. Frozen soil (5 g) was thawed and suspended in lysis buffer and boiled for 15 minutes as described previously (42). The supernatant was retained and amended with 100% chilled TCA to final concentration of 25% (vol/vol) and kept at −20°C overnight. Samples were centrifuged at 21,000 x *g* for 20 min and the protein pellets processed as described previously (43) and solubilized in 6 M guanidine buffer (6 M guanidine; 10 mM dithiothreitol [DTT] in Tris-CaCl_2_ buffer (10 mM Tris;, pH 7.8) with 3 hr incubation at 60°C. An aliquot of 25 µl/sample was retained for protein estimation and the rest of the protein sample was digested, peptides desalted and solvent exchanged as described earlier (44). The amount of protein extracted from each sample was calculated using the RC/DC protein estimation kit (Bio-Rad Laboratories, Hercules, CA, USA) as per the manufacturer’s instructions. Bovine serum albumin (supplied with the kit) was used as standard for the assay.

All chemicals were obtained from Sigma Chemical Co. (St. Louis, MO), unless specified otherwise. High performance liquid chromatography- (HPLC-) grade water and other solvents were obtained from Burdick & Jackson (Muskegon, MI), 99% formic acid was purchased from EM Science (Darmstadt, Germany) and sequencing-grade trypsin was acquired from Promega (Madison, WI).

### NanoLC-MS/MS Analysis

Peptides (75 ug) were loaded onto in-house prepared biphasic resin packed column [SCX (Luna, Phenomenex, Torrance, CA) and C18 (Aqua, Phenomenex, Torrance, CA)] as described earlier (44, 45) and subjected to an offline wash for 15 min as previously described (46). The sample column was aligned with an in-house C18 packed nanospray tip (New Objective, Woburn, MA) connected to a Proxeon (Odense, Denmark) nanospray source as previously detailed (46). Peptides were eluted and subjected to chromatographic separation and measurements via 24-hr Multi-Dimensional Protein Identification Technology (MuDPIT) approach as described earlier (44-46). Measurements were carried out using LTQ mass spectrometer (Thermo Fisher Scientific, Germany) coupled to the Ultimate 3000 HPLC system (Dionex, USA) and operated in data dependent mode, via Thermo Xcalibur software V2.1.0 as described earlier (45).

For protein identification, the raw spectra from each run were searched against a custom database and was constructed using protein sequences predicted from metagenome assemblies obtained from the same soil and 20-30 cm depth (14), metagenome assemblies from incubations (Supplementary Table 2), and reference proteomes for 47 common soil organisms (Supplementary Table 5). These predicted proteins were used for constructing a database for metaproteomic searches (available through http://enveomics.ce.gatech.edu/data/multiomics-soil). Database matching was done via Myrimatch v2.1 algorithm (47) set to parameters described before (48) with minor modifications where static cysteine and dynamic oxidation modifications were not considered. Identification of at least two peptides per protein (one unique and one non-unique) sequence was a prerequisite for protein identifications. Common contaminant peptide sequences from trypsin and keratin were concatenated to the database. Reverse database sequences were also included in the database as decoy sequences to calculate false discovery rate (FDR). For data analysis, spectral counts of identified peptides was normalized as described before (49) to obtain the normalized spectral abundance factor (NSAF) and the NSAF values were multiplied by a constant number (100,000) for better visualization and referred to as normalized spectral counts (nSpc). The nSpc were used to compare expression of proteins across different samples and different time points. Detected proteins predicted from metagenomic assemblies were annotated using BLASTp (50) and UniProt database as reference (51) (downloaded in May of 2017).

### Accession numbers

Raw metagenomic and metatranscriptomic soil datasets and MAGs are deposited in the European Nucleotide archive under study number PRJEB27434.

## ACKNOWLEDGMENTS

We thank Joel Kostka and Alissa Hooker for helpful discussions related to the manuscript. This work was supported in part by U.S. Department of Energy, Office of Biological and Environmental Research, Genomic Science Program [award DE-SC0006662], US National Science Foundation [Award 1831582], and the Chilean Fulbright-Conicyt doctoral scholarship [L.H.O.].

